# Pop-in: the inversion of pop-out for a feature dimension during visual search in area V4 of the monkey cortex

**DOI:** 10.1101/2022.06.23.497353

**Authors:** P. Christiaan Klink, Rob R.M. Teeuwen, Jeannette A.M. Lorteije, Pieter R. Roelfsema

**Affiliations:** Dept. Vision & Cognition, Netherlands Institute for Neuroscience, Royal Netherlands Academy of Arts & Sciences, Amsterdam, The Netherlands; Experimental Psychology, Helmholtz Institute, Utrecht University, Utrecht, The Netherlands; Laboratory of Visual Brain Therapy, Sorbonne Université, Institut National de la Santé et de la Recherche Médicale, Centre National de la Recherche Scientifique, Institut de la Vision, Paris F-75012, France; Dept. of Integrative Neurophysiology, Centre for Neurogenomics and Cognitive Research, VU University, Amsterdam, The Netherlands; Dept. of Psychiatry, Academic Medical Center, University of Amsterdam. Amsterdam, The Netherlands

**Author notes:** **Correspondence:** P.C. Klink, P.R. Roelfsema. Authors contributed equally (P. Christiaan Klink, Rob R.M. Teeuwen). **Competing Interest Statement:** No competing interests. **Author contributions (CrediT):** P.C. Klink: Conceptualization, Methodology, Software, Investigation, Formal Analysis, Resources, Data Curation, Writing – Original Draft Preparation, Writing – Review & Editing, Visualization, Project Administration, Funding Acquisition; R.R.M. Teeuwen: Formal Analysis, Data Curation, Writing – Original Draft Preparation, Writing – Review & Editing, Visualization; J.A.M. Lorteije: Conceptualization, Investigation, Writing – Review & Editing; P.R. Roelfsema: Conceptualization, Resources, Supervision, Funding Acquisition, Writing – Review & Editing. **Data deposition:** All Data & Analysis Code reported in this paper are available on GIN (https://gin.g-node.org/ChrisKlink/NHP_VisualSearch_Pop-in). A doi will be associated with the dataset upon acceptance of the final version of this manuscript.

**Keywords:** Visual search, V4, monkey, suppression, enhancement

## Abstract

During visual search, it is important to reduce the interference of distracting objects in the scene. The neuronal responses elicited by the search target stimulus are typically enhanced. However, it is equally important to suppress the representations of distracting stimuli, especially if they are salient and capture attention. We trained monkeys to make an eye movement to a unique ‘pop-out’ shape stimulus among an array of distracting stimuli. One of these distractors had a salient color that varied across trials and differed from the color of the other stimuli, causing it to also pop-out. The monkeys were able to select the pop-out shape target with high accuracy and actively avoided the pop-out color distractor. This behavioral pattern was reflected in the activity of neurons in area V4. Responses to the shape targets were enhanced, while the activity evoked by the pop-out color distractor was only briefly enhanced, directly followed by a sustained period of pronounced suppression. These behavioral and neuronal results demonstrate a cortical selection mechanism that rapidly inverts a pop-out signal to ‘pop-in’ for an entire feature dimension thereby facilitating goal-directed visual search in the presence of salient distractors.

**Significance statement:** Goal-directed behaviors like visual search involve both the selection of behaviorally relevant targets and the suppression of task-irrelevant distractors. This is especially important if distractors are salient and capture attention. Here we demonstrate that non-human primates suppress a salient color distractor while searching for a target that is defined by shape, i.e. another feature dimension. The neuronal activity of V4 neurons revealed the temporal evolution of target selection and distractor suppression. The neuronal responses elicited by the pop-out target stimuli were enhanced whereas responses elicited by salient pop-out color distractors were suppressed, after an initial brief phase of response enhancement. Our results reveal a ‘pop-in’ mechanism by which the visual cortex inverts an attentional capture signal into suppression to facilitate visual search.

## Introduction

Humans and animals usually need to select one of several stimuli for action. This selection process relies on priority signals in the brain such as the salience of stimuli and the subject’s goals (1–7). In the visual domain, for example, one could be faced with the task of locating a target object among distractor objects, e.g., trying to find one’s keys on a cluttered desk (Fig. 1A). A combination of bottom-up and top-down processes often solves this problem (3). If the keys have a high saliency because they are bright red, for example, they ‘pop out’ from the background, which would be considered a bottom-up contribution. However, top-down factors also play an important role. You may, for example, imagine the shape of your keychain or try to remember where the keys most likely are. Visual search is therefore a very useful experimental paradigm to study the role of bottom-up and top-down factors in visual selection.

**Figure 1.**
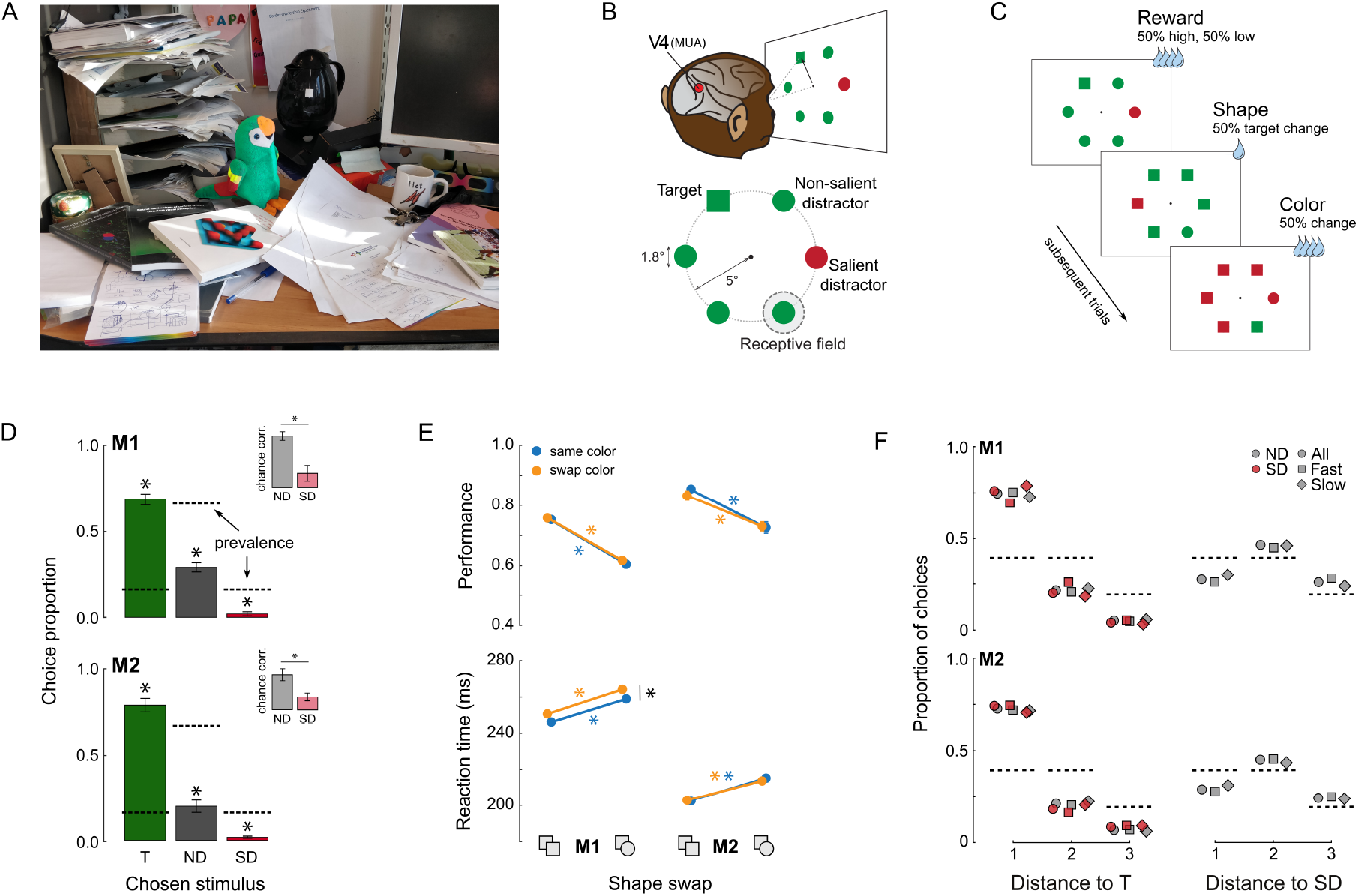
Task description and behavioral results. **A)** Real-life example of visual search with a salient distractor. When looking for your keys on a crowded desk, you may be looking for small key-shaped objects. Your attention may however be captured by salient objects like the bright green parrot, which might interfere with the process of finding your keys. **B)** We recorded from area V4 while monkeys performed a visual search task in which they selected the odd-shape-out (here a square among circles) with an eye movement. One of the six visual items was in the V4 receptive field. The target was the stimulus that differed from the others by shape. Non-salient distractor stimuli had the same color as the target, while a single salient distractor stimulus popped out because it had a different color. **C)** Example series of three trials. In the second trial the target and distractor shapes swapped with respect to the first trial (this occurred 50% of the time). In the third trial, the target and distractor colors swapped (this also occurred 50% of the time). In addition, the reward magnitude was randomly varied (50% high, 50% low). **D)** Accuracy (green bars) and the proportion of trials on which the monkeys made an error by choosing a non-salient distractor (ND, grey bars) or the salient distractor (SD, red bars). Non-salient distractors are 4 times more prevalent than targets and salient distractors (prevalence indicated with dashed horizontal lines). The insets show the proportion of choices of distractor stimuli corrected for prevalence. Even after this correction, the animals chose the salient distractor less often than the non-salient distractors (* indicates p < 0.001 for a one-tailed t-test SD < ND). Error bars indicate the standard deviation over recording sessions. **E)** The effects of color and shape swaps on accuracy (top panel) and reaction time (bottom panel) for both monkeys. Yellow lines indicate trials in which the target and salient distractor colors swapped relative to the previous trial; blue lines are trials in which those colors stayed the same. The horizontal axis indicates whether the target shape changed relative to the previous trial. Error bars (often smaller than the data points) indicate S.E.M., asterisks denote p < 0.001 for main effects as indicated by two-way ANOVAs (no interaction effects were significant at p < 0.05). **F)** Dependence of erroneous choices on the relative locations of the target (T) or salient distractor (SD) stimuli. The proportion of choices on error trials is plotted as function of the distance between the chosen stimulus in the search array (a distance of one indicates the two stimuli were next to each other, a distance of two means there was one stimulus in between, etc.), the identity of the chosen stimulus (grey: ND; red: SD), and the reaction time (30% fastest and slowest response indicated with square and diamond symbols respectively). The dashed lines indicate chance level.

In many bottom-up search paradigms, the target pops out, because it has a unique feature. For example, it is the only bright red item among grey distractors, or it is the only circle in the display in which all other elements are squares. There are versions of this paradigm in which subjects do not know beforehand what they will be looking for, but only that it is the unique item. For example, the display might have either one square among circle distractors or one circle among square distractors. The search for items with unique properties is usually parallel, which means that the time to find an item does not depend strongly on the total number of distractors in a search display (7). Previous studies on the neuronal correlates of pop-out search demonstrated that the responses elicited by pop-out stimuli are stronger in the visual, parietal, and frontal cortex than the responses to stimuli that do not pop-out (8–18). In top-down search paradigms, the subject looks for a specific item known as ‘search-template’ (21, 22). The search template represents a top-down influence on visual selection (1, 23) and the representations of the items in the display that match the search template are also enhanced in areas of the visual, parietal, and frontal cortex (19, 22–29).

Many displays contain salient distractors that interfere with visual search. This is the case in Figure 1 for the green parrot, which captures attention, making it more difficult to find the keys. Researchers have debated the degree of automaticity of attentional capture, with some researchers arguing that it is mandatory (30) whereas others arguing that it can be prevented by sufficiently strong top-down signals (31). Importantly, conditions exist under which salient display items do not appear to interfere with visual search (32, 33) or cause even less interference than regular, non-salient distractors (34–37).

The mechanism by which salient distractors can be suppressed is not yet fully understood and there are contrasting views (38). One possibility is that salient distractors initially capture attention, but that it is rapidly curtailed by top-down suppression mechanisms (39). Support for such reactive suppression comes from human EEG studies employing markers of distractor selection and suppression (40–43). The signal suppression hypothesis (35–37) proposed another account, in which a top-down influence prevents the capture of attention by salient distractors so that there is no need for disengagement. This viewpoint received support from behavioral studies (34, 35, 44) and other human EEG studies (34, 36, 37, 45, 46). We note, however, that the relation between this putative suppressive signal and its EEG signatures is under dispute (43, 47).

The degree to which salient distractors attract attention and, hence the need for disengagement, depends on how predictable they are. Salient distractors are more efficiently suppressed if their features are predictable, for example, because they are the same across trials or are known in advance (48, 49). Bichot et al. (50) demonstrated that the representations of stimuli that consistently appear as distractors, across many days, are strongly suppressed in the frontal cortex of monkeys. Like distractor predictability, foreknowledge about the target also decreases the influence of salient distractors. If the subject knows the target, a search template can be established before the display appears and the influence of salient distractors is weaker than in pop-out search in which the target properties are not specified. Researchers proposed that pop-out search demands a special ‘singleton detection mode’ (32). If subjects search for a salient target with unknown features, they are more susceptible for interference by salient distractors. The degree of interference by the distractor depends on the relation between the features of the target and the distractor (35, 36, 38, 51, 52). Interference is strong if the target and salient distractor are defined on the same feature dimension, e.g., if they both have an orientation that differs from that of all other distractors. Interference is weaker if they are defined on a different feature dimension, e.g., the target differs in orientation from the other items whereas the salient distractor differs in color. In this situation, the features can be weighted. The target dimension receives a higher weight than the salient distractor so that the degree of distraction can be diminished (40, 52–54).

Two previous studies have examined the neuronal mechanisms for the suppression of salient distractors during visual search. Ipata et al. (55) had monkeys searching for a black target shape among black distractors. They added a salient distractor, which was green and bright, and recorded neurons in the lateral intraparietal area (LIP) of the parietal cortex. As expected, targets elicited stronger neuronal responses than the black distractors, but the activity elicited by the salient green items was even weaker than that elicited by the regular black distractors. Hence, the representation of the salient distractor is efficiently suppressed in the parietal cortex. A later study by Cosman et al. (56) replicated this finding in the frontal eye fields (FEF) in a task where the monkeys searched for a white target letter while the salient distractor was colored. Again, the target letter elicited strongest activity, followed by the regular distractors and the salient distractor elicited weakest activity. These results are in accordance with those of Bichot et al. (50) showing the effective suppression of a specific feature that is always distracting in the frontal cortex. (57, 58). However, parietal and frontal cortex are relatively high up in the cortical processing hierarchy and activity elicited by salient distractors might still be enhanced in the visual cortex, even after extensive training. The representation of salient distractors in visual cortex remains to be investigated.

In the present study we tested the generality of the suppression mechanisms by asking three questions: (1) Are salient distractors suppressed in the visual cortex? (2) Is the efficient suppression of a salient distractor stimulus also possible when its features vary across trials? (3) Can salient distractor suppression occur when the subject searches for a pop-out stimulus on a different feature dimension?

We trained monkeys in a task in which they carried out a pop-out search for a shape while we presented a salient color distractor with a color that varied across trials. They had to select the shape singleton as target for an eye movement to obtain a juice reward. As expected, the shape singleton elicited stronger V4 activity than the distractors with a different shape. Remarkably, the V4 representation of salient color singleton was briefly enhanced followed by a period of pronounced suppression below the level of representation of the regular non-pop-out distractors, even though its color was unpredictable. At a behavioral level, the monkeys also selected the salient distractor less often than the regular distractors, indicating active avoidance. We conclude that after extensive training, the neuronal mechanisms for visual search can exploit the presence of a color singleton if it is always a distractor, and rapidly cause it to ‘pop-in’ instead of pop-out, thus avoiding capture and promoting efficient goal-directed behavior.

## Results

Two monkeys were extensively trained to perform a visual search task (Fig. 1B,C) in which they had to select a single odd-shape-out (target) from an array of six stimuli. On any given trial, the target could either be a circle among squares, or a square among circles. To study whether V4 neurons show suppression of salient distractors, one of the distractor stimuli had a different color than the others (either green among red, or red among green) (Fig. 1B, bottom). The shapes, colors, and locations of the target and salient distractor were randomly assigned on each trial so that the animal could not predict the shape or color of the target and salient distractor. As a result, consecutive trials could have the same shape and colors assigned to the target and distractor, both could change, or one of the feature assignments could stay the same while the other changed. Moreover, to examine a previously reported interaction between stimulus salience and reward in human visual search behavior (59), we randomly rewarded correct responses with either small or large juice rewards (with the large reward being approximately four times the small reward amount). After an initial training phase to learn the task, both monkeys were extensively trained to reach high performance levels (22 training sessions for M1, 56 for M2).

We recorded 34,543 trials in monkey 1 (M1) and 13,815 trials in monkey 2 (M2) in 28 and 16 sessions, respectively. Both monkeys displayed similar eye movement patterns (Fig. 1D), most often choosing the target stimulus (M1: 69%, M2: 78% of choices), followed by non-salient distractors (M1: 29%, M2: 20%), and only rarely choosing the salient distractor stimulus (M1: 2%, M2: 2%). The lower probability of choosing a salient distractor than a non-salient distractor remained when we accounted for the fact that there were four non-salient distractors and only one salient distractor (see Fig. 1D insets, corrected for prevalence). The probability of choosing the target was much higher than chance (one-tailed t-test, M1: t(26) = 87.4, p < 0.001; M2: t(15) = 53.5, p < 0.001). On error trials, both animals were significantly less likely to choose the salient distractor than a non-salient distractor (prevalence-corrected, one-tailed paired t-test, M1: t(26) = - 51.6, p < 0.001; M2: t(15) = -21.6, p < 0.001).

Swapping the colors of the target and salient distractor on successive trials did not affect accuracy for either animal as indicated by a two-way ANOVA with color-swap and reward quantity as independent variables (all ps > 0.48). It did slow down M1 by a few milliseconds (Fig. 1E; F(1, 12210) = 38.8, p < 0.001), but had no effect on M2’s reaction time (F(1,7484) = 1.59, p = 0.83). A change of the target shape had a much more pronounced effect of performance. It decreased the accuracy of both animals and increased the reaction times (Fig. 1E; all p < 0.001). There were no interactions between the effects of color and shape changes. These results imply a shape-based priming of pop-out effect across trials (60), but an absence of color-based priming, which is consistent with the animals being in ‘shape-searching’ mode due to extensive training on the ‘odd-shape-out’ search task. Unlike previous work in humans (59), we did not observe any main or interaction effects of reward quantity on visual search performance (Supplemental Fig. 1).

What happened when the monkeys made an error? They predominantly selected the distractor stimulus that was adjacent to the target in the search array (Fig. 1F), a pattern that was neither influenced by the location of the salient distractor, nor by the saccadic reaction time (comparing the 30% fastest and 30% slowest saccades) (squares and diamonds in Fig. 1F). The distribution of erroneous saccades relative to the target position was the same for salient and non-salient distractors (red and grey symbols in Fig. 1F), which indicates that the probability of choosing the salient distractors was decreased uniformly (Fig. 1D) with little influence of the target location.

Whereas the signal suppression hypothesis (34) proposes that a salient distractor can be proactively suppressed to avoid attentional capture, the stimulus-driven rapid-disengagement account suggests that capture does temporarily occur but that it is then quickly suppressed. The latter scenario should be associated with a brief period of pop-out for the salient distractor followed by a sustained period of distractor suppression. Because visually guided saccades can occur at very low latencies in both humans and monkeys (61–65), especially after prolonged training (66), we wondered whether an early neuronal pop-out of the salient distractor would result in very rapid saccadic responses to the salient distractor before the distractor suppression could have manifested. To investigate this possibility, we compared the distributions of saccade reaction times (SRTs, Supplemental Fig. 2) for target and salient distractor choices. A larger proportion of the salient distractor choices than the target choices occurred at the shortest reaction times in both monkeys (Fig. 2A). We calculated the proportion of salient distractor choices (*p*_*SD*_ *= n*_*SD*_*/n*_*ALL*_) as function of SRT (Fig. 2B). In both animals, the proportion SD choices was significantly higher for the 12.5% shortest SRTs (first octile) than for SRTs in the 2^nd^-4^th^ octiles (chi-squared test, M1: X^2^(1) = 8.55, p < 0.01; M2: X^2^(1) = 21.41, p < 0.001). In M1 there was even a brief epoch in which the salient distractor was chosen more often than the target, but saccades to the salient distractor were strongly suppressed for longer SRTs. Also, in M2 the salient distractor choices decreased for longer SRTs, but the target was always chosen with the highest probability (Supplemental Fig. 2). This result indicates that the distractor pops out in an early interval after stimulus presentation, but that the pop-out signal is rapidly suppressed to prevent erroneous choices.

**Figure 2.**
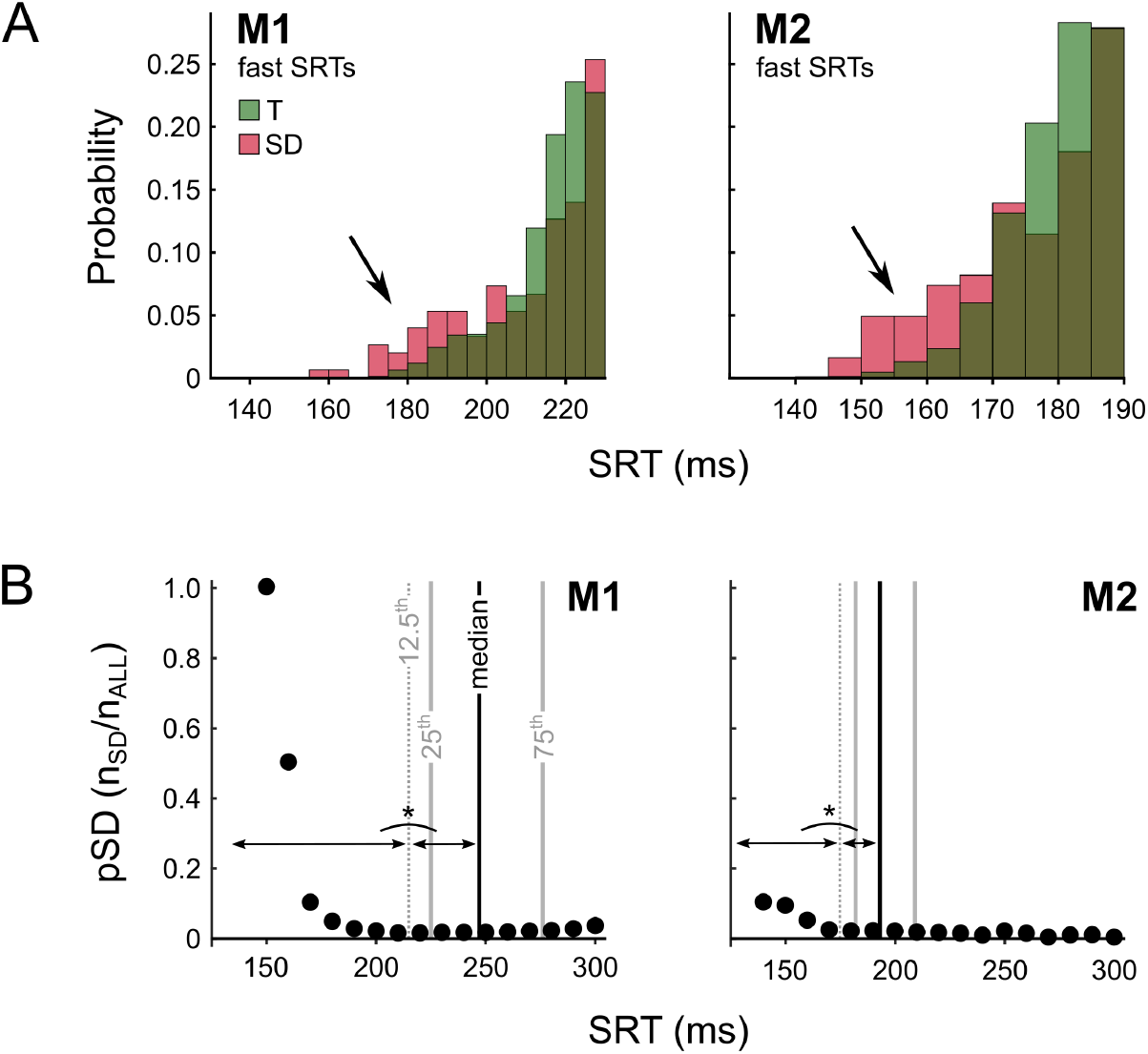
Saccadic reaction times and choices. **A)** Distributions of shortest saccadic reaction times (SRTs, fastest 25^th^ percentile) for target (T, green) and salient distractor choices (SD, red) in the two monkeys. The distributions were normalized such that both the red and green bars sum up to 100% (see Supplemental Fig. 2 for the full SRT distributions, normalized within choice type (as here) and also by to the total number of saccades). The dark colors indicate overlap between the red and green distributions. The probability of choosing the salient distractor was increased at short SRTs (black arrows). **B)** Proportion of salient distractor choices (*pSD*) calculated in a sliding 20 ms window, moving at 10 ms increments. Solid vertical lines are the median, 25^th^, and 75^th^ percentiles of the full SRT distributions. In both monkeys, the proportion of salient distractor choices is significantly higher for the 12.5% fastest responses (first octile, left of the dashed vertical line) than in the second through fourth octiles (chi-squared test, M1: X^2^(1) = 8.55, p < 0.01; M2: X^2^(1) = 21.41, p < 0.001).

Next, we compared the neuronal responses in V4 elicited by target stimuli, non-salient distractor stimuli and salient distractor stimuli on correct trials (Fig. 3A, top panels). We pooled the data across animals (Fig. 3, left panels) because the results were similar for M1 and M2 (Fig. 3, middle and right panels). The late V4 response elicited by target stimuli was stronger than that elicited by non-salient distractor stimuli (time window 150-200 ms after stimulus onset, t(34) = 8.9, p < 0.001; M1: t(9) = 5.6, p < 0.001; M2: t(24) = 7.0, p < 0.001). The response elicited by the salient distractor stimulus was weaker than that elicited by the target stimulus and, importantly, also weaker than that elicited by the non-salient distractor stimulus (t(34) = -9.9, p < 0.001; M1: t(9) = -5.4, p < 0.001; M2: t(24) = -9.1, p < 0.001). This ordering of response strength was very consistent among recording sites (Supplemental Fig. 3).

**Figure 3.**
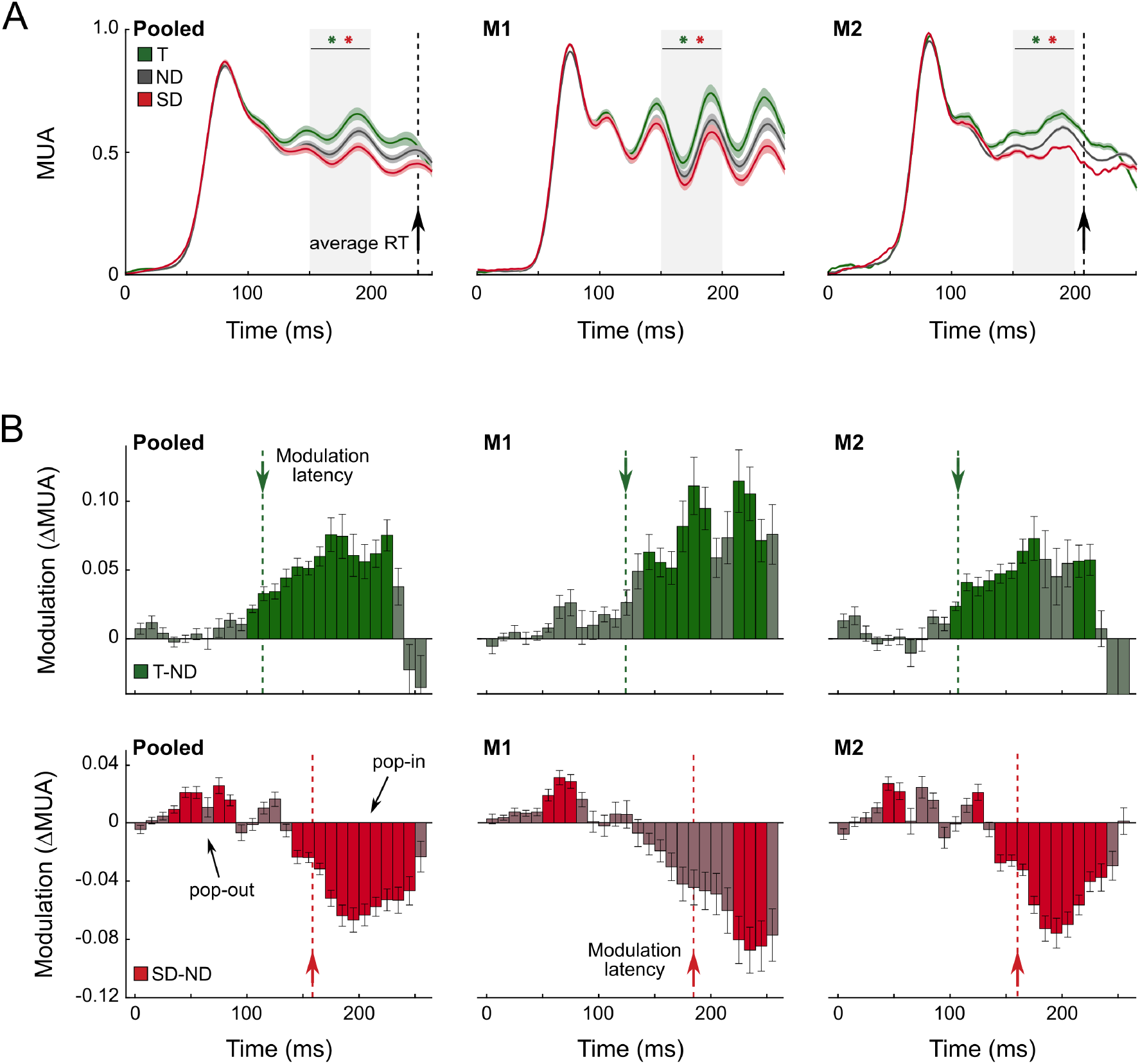
V4 activity during visual search reveals the time-course of pop-out and pop-in. **A)** Neuronal responses in area V4 responses on correct trials. Average V4 activity elicited by the target (T, green trace), non-salient distractors (ND, gray trace) and the salient distractor (SD, red trace) averaged across animals (left panel) and for individual monkeys (M1: middle panel; M2: right panel). Shaded area corresponds to S.E.M. across recording sites. Black arrows indicate the average reaction time (for M1 this was later than 250 ms and is not depicted). The light grey areas indicate the time window used for statistical testing of the response modulation, with * indicating p < 0.001 with a paired t-test (green: T-ND; red: SD-ND). **B)** Time-course of neuronal target and salient distractor modulation. Top row, difference in activity elicited by the target and non-salient distractor (T-ND; non-overlapping 10 ms time bins) pooled across monkeys (left) and individual animals (middle and right panels). Green bars indicate significant epochs at p < 0.05 (t-test with Bonferroni correction for multiple comparisons). Bottom row, difference in activity elicited by the salient distractor and non-salient distractor (SD-ND) with the red bars indicating p < 0.05 (t-test, Bonferroni correction). In both animals, there is an initial epoch of salient distractor enhancement, followed by suppression, later than 150 ms. Colored arrows indicate the latency of target enhancement (green) and salient distractor suppression (red).

We examined the time-course of target enhancement and salient distractor suppression by subtracting V4 activity elicited by the non-salient distractor stimuli from the other two conditions (Fig. 3A, bottom panels). We measured the latency of the enhancement and suppression of targets and salient distractors with a fitting procedure that has been described before (67) (see Methods and Supplemental Fig. 4). The latency of target enhancement was 112 ± 9 ms (averaged across monkeys, standard deviation determined with bootstrapping) and the latency of suppression of salient distractors was 158 ± 25 ms. This pattern was also present in individual animals (M1_T_: 124 ± 20 ms, M2_T_: 108 ± 15 ms; M1_SD_: 184 ± 14 ms, M2_SD_: 159 ± 11 ms) and the salient distractor suppression was significantly later than the target enhancement (paired t-test, M1: t(75) = - 26.3, p < 0.001; M2: t(72) = -22.1, p < 0.001; Pooled data: t(67) = -17.7, p < 0.001). Thus, the pop-in effect was expressed in area V4 as a decreased response to the irrelevant singleton, even though its color was unpredictable.

The brief early epoch with an enhanced probability of saccades to the salient distractor suggests that the distractor representation might be briefly enhanced in V4 (33, 35, 37) before it is suppressed. We therefore examined the possibility of an early response enhancement. We observed that the salient distractor (Fig. 3B, red bars) indeed caused a brief epoch of enhanced activity before suppression became evident, in a time-window up to 100 ms after stimulus onset (Fig. 3B shows significant modulation in several 10 ms non-overlapping time bins in both monkeys; t-tests at p < 0.05, Bonferroni corrected).

We also examined a possible influence of the behavioral priming effect, which occurred when the target shape was the same on consecutive trials, on V4 activity. The priming effect did not have a consistent influence on V4 activity (Supplemental Fig. 5), which suggests that the increase in SRT may originate in downstream brain regions, as a post-selective process (54). Furthermore, V4 activity on error trials was more variable than on correct trials (Supplemental Fig. 3).

## Discussion

Goal-directed behaviors require a selection process that highlights relevant stimuli and suppresses distractors. Here, we used a visual search paradigm to investigate the representations of relevant and irrelevant pop-out stimuli (7) in area V4 of the monkey visual cortex. We presented a salient pop-out color distractor with an unpredictable color while the monkeys searched for a singleton shape. Our results demonstrate that the visual brain can suppress the representation of pop-out stimuli on an irrelevant feature dimension while enhancing the representation of pop-out stimuli on a relevant feature dimension. A brief neuronal activity enhancement preceded the suppression of distractor representations (Fig. 4), suggesting that an initial pop-out process is required before it can invert into pop-in. To our knowledge, this is the first demonstration of ‘pop-in’ for an irrelevant feature dimension, which presumably emerged during the monkeys’ considerable training.

**Figure 4.**
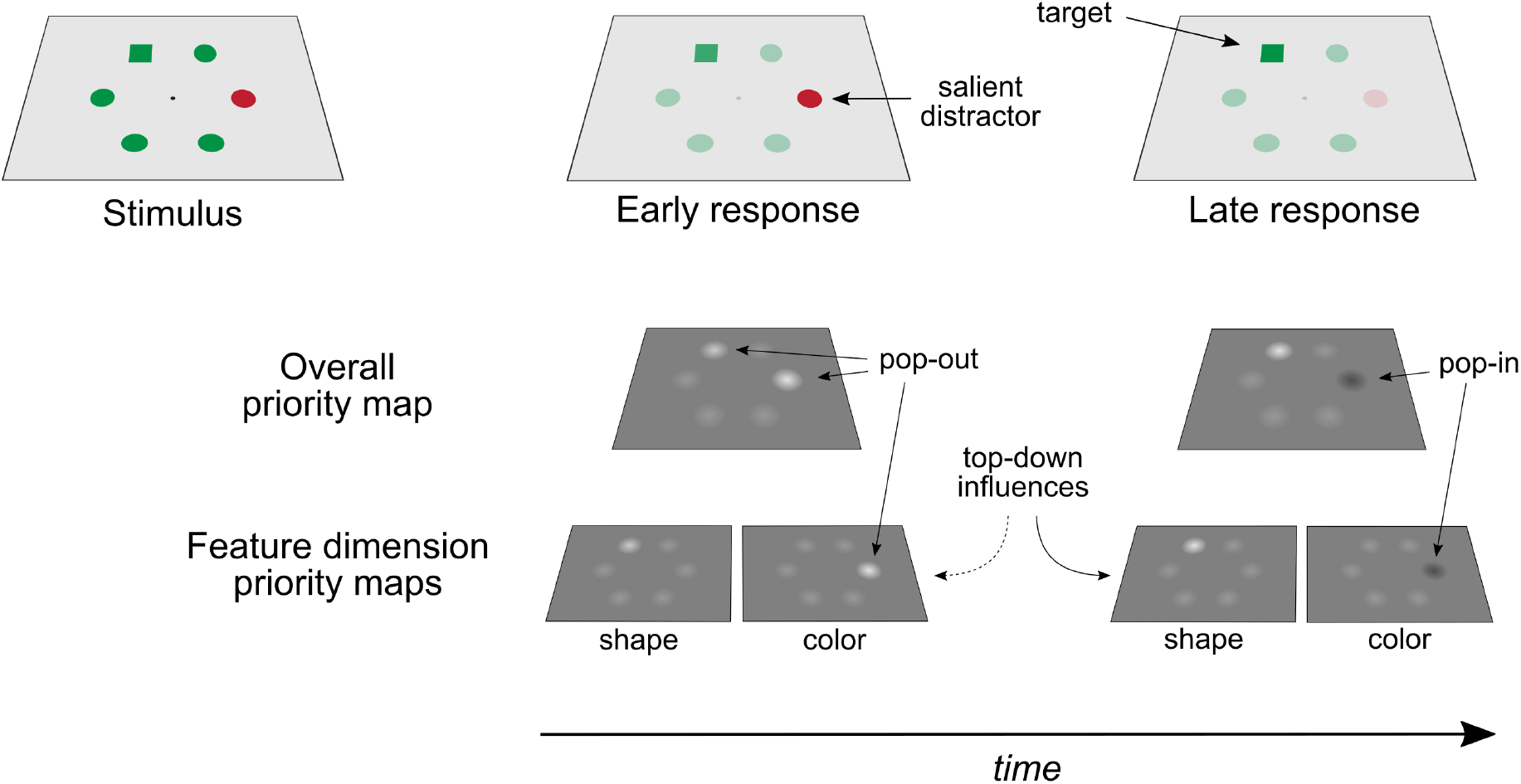
Pop-out and pop-in. During the early phase of the V4 response (middle) to a visual search stimulus (left), both the shape and color singletons pop-out. In a later phase of the response (right), top-down influences invert the pop-out of the salient color distractor into pop-in.

The efficiency of visual search depends on bottom-up factors that determine the salience of stimuli, such as brightness and local feature contrasts causing pop-out, and the top-down search template, the internal representation of the item that the subject is searching for (1, 3, 4, 22). Researchers have hypothesized that stimulus salience and goal-driven influences on the distribution of attention jointly determine a ‘priority map’ of visual space (3–7, 68–70). There are multiple candidate brain regions for such a priority map, including the LGN (68), pulvinar (71), superior colliculus (72, 73), V1 (74), V4 (66), the parietal (8, 10, 55) and prefrontal cortex (11). Indeed, stimulus-driven pop-out signals have a widespread influence on the neuronal firing rates in early visual cortex (12, 75, 76), parietal cortex (10), frontal cortex (11), and subcortical structures like the superior colliculus (77). Similarly, the top-down influences of the search template on firing rates also occur in most, if not all, of the same brain regions, including V1 (78, 79), V4 (15, 18), the parietal (55) and prefrontal cortex (11, 50, 56). It is conceivable that the relative contributions of the multiple priority maps depend on the task, e.g., on the features that matter and on whether the subject reports the location of the target with an eye or hand movement.

There are many instances in which the representation of visually salient items needs to be suppressed, because task relevant items are less conspicuous, causing a conflict between bottom-up and top-down factors. The signal suppression hypothesis (36, 37) proposed that top-down suppression signals can prevent attentional capture by salient distractors if their features are known in advance (34–37, 44–46, 51, 56, 80– 82). An alternative possibility is that salient distractors attract attention, but that it is rapidly disengaged (39). Previous electrophysiological studies in areas LIP and FEF of monkeys revealed that the neuronal activity elicited by a salient distractor with a predictable color can indeed be suppressed below the activity elicited by regular distractors (55, 56). The suppression of salient distractors has also been measured as a distractor positivity (Pd) component in the EEG of humans (37, 38, 83) but a recent study using the steady-state visually evoked potential (SSVEP) did not find evidence for suppression below the activity elicited by regular distractors (47). This SSVEP study used displays with only few items, however, and it has been suggested that such displays do not emphasize pop out but require other search processes (‘clump scanning’) (41, 52, 84). The present study went beyond these previous studies by investigating whether suppressive signals influence spiking activity in the visual cortex of monkeys. Furthermore, we used a new task in which the features of the salient distractor were unpredictable, and the monkey was searching for a pop-out stimulus on a different feature dimension.

Unlike the previous studies (55, 56), we found that the salient distractor elicited a brief enhancement of V4 activity that later inverted into sustained suppression. It seems likely that the early response enhancement occurred because the color of the salient distractor was unpredictable so that it first needed to be registered before it could be suppressed. We also observed a behavioral consequence of this brief pop-out phase because a proportion of the early saccades landed on the salient distractor whereas it was less likely to be selected than regular distractors at later time points, when pop-out has inverted into pop-in. This result provides evidence for short-lived attentional capture, followed by rapid attentional disengagement (38, 39). It supports previous EEG studies indicating that predictable salient distractors cause attentional capture (40) and is not in accordance with the signal suppression theory, which proposed that suppressive top-down signals eliminate attentional capture.

In a previous study on the role of area V4 during visual search Ogawa and Komatsu (15) trained monkeys to search for either shape or color singletons in displays that also included a singleton in the other dimension, as a salient distractor. Unlike in the present study, however, the monkeys searched for shape and color singletons in alternating blocks of trials. When they made an error, they selected the salient distractor more often than regular distractors, which is also different from the current results. Accordingly, V4 activity elicited by the target of search was strongest, V4 activity elicited by salient distractors was intermediate and stronger than that elicited by regular distractors (15). In other words, in the previous study both the singleton target and the singleton distractor popped out, while in the present study, the color singleton was never relevant and its V4 representation was suppressed below the activity elicited by regular distractors.

Our results indicate that there are at least two processing steps in the present search task (Fig. 4). Initially, there is pop-out in two feature dimensions: color and shape. Later in the trial, the activity elicited by the shape singleton remains enhanced, whereas activity elicited by the color singleton is suppressed, indicating that V4 could contribute to a priority map with enhanced target and suppressed distractor representations (Fig. 4). The source of the suppressive pop-in signal is unknown, but it could rely on feedback projections (85, 86) that might have been strengthened during training. In accordance with this view, microstimulation of FEF interacts with stimulus driven activity in early visual cortex in a topographic manner, with an effect that depends on stimulus strength and the presence of distractors (87). It is remarkable that the neuronal mechanisms for the registration of the salient distractor and its later suppression can co-exist with the mechanisms for pop-out on another feature dimension. Previous studies anticipated that that the degree to which different feature dimensions cause pop-out can be weighted (52–54). However, to our knowledge, these theories did not anticipate that dimension weights could become negative, causing attentional repulsion of singletons on a specific feature dimension.

Previous studies demonstrated a profound influence of the recent history of trial types during visual search. Repeatedly searching for the same stimulus features causes priming. It reduces an observer’s reaction time, improves accuracy, and increases the difference between the strengths of the neuronal representation of targets and distractors (9, 18, 60, 88–92). We here observed a priming effect of shape. When the shape of the search target remained the same on consecutive trials, the monkeys were faster and more accurate than when it was different. Interestingly, we did not find a behavioral priming effect of color as was observed in previous studies (9, 18, 88), in which the search target was a color singleton. It therefore seems likely that priming only occurs for the feature dimension that defines the search goal.

Earlier studies also demonstrated an effect of reward quantity on visual search performance (59, 93–95). A study in human subjects demonstrated that visual search is faster if a preceding trial with the same target and distractor features gave rise to high, as opposed to low, reward (59). We did not replicate this effect in our monkeys, where reward magnitude on the previous trial did not strongly affect performance. One possible explanation is that the animals were highly trained, which may have reduced their sensitivity to reward outcomes on individual trials. However, other explanations, including species differences, are also conceivable.

In conclusion, our work shows parallel mechanisms of target enhancement and salient distractor suppression during visual search in V4 that rapidly develop and manifest behaviorally as efficient distractor avoidance and goal-directed target selection. It seems likely that the conversion of initial distractor enhancement into functional and profound that occurs round 150 ms after stimulus onset reflects a top-down dynamic adjustment of the weights of individual feature dimensions. The extended training history, during which the salient color never coincided with the search target, must have engaged plasticity mechanisms inverting pop-out into pop-in, making the mechanisms of visual search more versatile than might have been anticipated.

## Acknowledgements

We thank Kor Brandsma, Anneke Ditewig, and Lex Beekman for animal care and biotechnical assistance; Bram van Vugt and Pia Jentgens for assistance with data acquisition and animal training; Matthew Self for graciously allowing us to photograph his desk for the scene depicted in Figure 1; and Jan Theeuwes and Daniël Schreij for early discussions of the work. The in-house experimental control software was originally developed by Chris van der Togt. This work was supported by NWO (Crossover Program 17619 “INTENSE”; VENI 451.13.023), the European Union FP7 (ERC 339490 “Cortic_al_gorithms”), the Human Brain Project (Agreement No. 945539, “Human Brain Project SGA3”), and the Friends Foundation of the Netherlands Institute for Neuroscience.

## Methods

### Subjects

All animal procedures complied with the NIH Guide for Care and Use of Laboratory Animals, and were approved by the institutional animal care and use committee of the Royal Netherlands Academy of Arts and Sciences. Two male macaque monkeys participated in the experiment. They were 5 (M1) and 8 (M2) years old at the start of the experiments and weighted between 7-8 (M1) and 8-9 (M2) kg over the course of the recordings. The monkeys were socially housed in pairs in a specialized primate facility with natural daylight, controlled humidity and temperature. The home-cage was a large floor-to-ceiling cage that allowed natural climbing and swinging behavior. The cage had a solid floor, covered with sawdust, and was enriched with toys and foraging items. The diet consisted of monkey chow supplemented with fresh fruit. The access to fluid was controlled, according to a carefully designed regime for fluid uptake. During weekdays the animals received diluted fruit juice in the experimental set-up upon correctly performed trials. We ensured that the animals drank sufficient fluid in the set-up and supplemented extra fluid after the recording session if the monkeys did not drink enough. In the weekend the animals received at least 700 ml of water in the home-cage supplied in a drinking bottle. The animals were regularly checked by veterinary staff and animal caretakers and their weight and general appearance were recorded in an electronic logbook daily during fluid-control periods.

### Surgical procedures and training

We implanted both monkeys with a titanium head-post (Crist instruments) under aseptic conditions and general anesthesia as reported previously (96–98). The monkeys were first trained to fixate a 0.5 diameter fixation dot and hold their eyes within a small fixation window (1.2 diameter). They then underwent a second operation to implant arrays of 4x4, 4x5 and 5x5 micro-electrodes (Blackrock Microsystems) in V4. The inter-electrode spacing of the arrays was 400 µm. The animals were later extensively trained to perform the visual search task at adequate performance levels (22 training sessions with the final task for M1, 56 sessions for M2). During the early phase of the training the animals were required to make an eye movement from the fixation point to a single target, and in later phases the distractors were introduced at low contrast which over sessions gradually increased to the same contrast as the target.

### Electrophysiology

Recordings from the chronically implanted electrode arrays were made with TDT (Tucker Davis Technology) recording equipment using a high-impedance head-stage (RA16AC) and a preamplifier (either RA16SD or PZ2). The signal was referenced to a subdural electrode and digitized at 24.4 kHz. It was band-pass filtered (2^nd^ order Butterworth filter, 500 Hz – 5 kHz) to isolate high-frequency (spiking) activity. This signal was rectified (negative becomes positive) and low-pass filtered (corner frequency = 200 Hz) to produce multi-unit activity (MUA), which is the envelope of the high-frequency activity (99). MUA reflects the spiking of neurons within 100-150 mm of the electrode and MUA population responses are very similar to those obtained by pooling across single units (98–102). We used a video-camera based eye-tracker (Thomas Recording) to measure the eye position at a sampling frequency of 250 Hz. V4 receptive fields were mapped by presenting white squares (1°, luminance 115 cd/m^2^) on a dark background (2 cd/m^2^) at different positions of a grid (1° spacing). We defined the RF borders as the locations where activity fell below 50% of the maximum (103).

We removed trials with artifacts first by calculating the time-average for each trial and removing trials with extreme average MUA responses. We used an iterative z-scoring procedure (values higher than 3 were removed). If z-scores higher than 20 remained in the cleaned collection of trials, the process was repeated, leading to the removal of less than 2% of all the trials. We also removed trials that included any samples (without averaging) that had a z-score higher than 10. To normalize MUA, we subtracted the spontaneous activity level in a 100 ms time window prior to the onset of the stimulus and divided by the peak response after LOWESS smoothing (26 ms window). We only included recording sites with a signal-to-noise (SNR) higher than 2.5. SNR was computed for individual recording sessions by dividing the peak of the smoothed response by the standard deviation of the spontaneous activity level across trials. We excluded recording sites with fewer than 3 recording sessions that met the SNR criterion. For the other recording sites, we averaged the activity per recording site across sessions so that every recording site contributed only once to the statistics.

### Behavioral task and stimuli

Stimuli were presented on a 21” CRT monitor (Dell Trinitron) with a refresh rate of 85 Hz and a resolution of 1024x768 pixels, viewed at a distance of 87 cm. All stimuli were created using the COGENT graphics toolbox (developed by John Romaya at the LON at the Wellcome Department of Imaging Neuroscience) running in MATLAB (Mathworks Inc.) with custom experimental control software (104). The monkeys were trained to perform a visual search task. A trial started when the monkey acquired fixation on a 0.3° red (26.2 cd/m^2^) fixation dot in the center of the screen. After 200 ms of fixation within a 1.2° diameter window, 6 stimuli appeared, arranged in a circle around the fixation point, at 5.3° eccentricity. Simultaneously, the fixation dot became green (98.6 cd/m^2^) cueing the monkey to make a saccade. The stimuli were visible for 2,000 ms, during which the monkey was required to respond. If the monkey failed to respond in time, the trial was classified as aborted. Each stimulus could be either a square or a circle and was either red (76.0 cd/m^2^) or green (114.1 cd/m^2^), presented on a gray background (54.2 cd/m^2^). Stimuli had a size of 1.8° diameter. On each trial, one stimulus had a different shape (the target stimulus), one stimulus had a different color (the salient distractor stimulus), and the 4 remaining stimuli (non-salient distractors) had the same color as the target stimulus and the same shape as the salient distractor. The task of the monkey was to make an eye movement to the target stimulus, while ignoring the salient and non-salient distractors. Choices were detected as the eye-position entering a 4° diameter circular window around one of the stimuli. Upon a correct response, the monkey received a juice reward. This reward was randomly selected to be either small or large (~4 times the small amount). The trials were ordered in a pseudorandom fashion. We recorded 34,543 trials across 28 sessions in monkey 1 and 13,815 trials across 16 sessions in monkey 2.

### Computation of target and salient distractor modulation

Average MUA responses for target, non-salient distractor, and salient distractor stimuli were calculated for individual monkeys and the pooled data. To compute target and salient distractor modulation we subtracted the response to non-salient distractors from the response to targets and salient distractors, respectively, for each recording site in a 150-200 ms time window after stimulus onset. As statistical test we used paired t-tests over recording sites. The time-courses of target and salient distractor modulation were furthermore evaluated by recalculating the modulation in 10 ms non-overlapping bins and statistically tested with a series of t-tests, using Bonferroni correction for multiple comparisons.

### Latency of target selection and distractor suppression

To estimate the latency of the enhancement of the representation of the target and the suppression of the representation of the salient distractor we used a fitting procedure that has been described before (67). Briefly, a cumulative gaussian function was fit to the difference between either the target and the non-salient distractor response (i.e., target modulation) or the non-salient distractor and the salient distractor response (i.e., salient distractor modulation). The latency is estimated as the time point at which the fit reaches 33% of its maximum (Supplemental Fig. 4). The fits were calculated based on the population responses, i.e., after averaging across recording sites. We used a bootstrapping procedure (100 times) with replacement to estimate the mean and standard deviation of these latency estimates and compared latencies of target and salient distractor modulations with paired t-tests.

### Saccadic reaction times

We investigated the susceptibility to attentional capture by the salient distractor as a function of saccadic reaction time (SRT). We removed SRTs that were faster than 75 ms because we deemed such responses to be too fast to be visually guided based on previous reports. This resulted in the removal of 6 target (M1: 2, M2: 4) and 9 salient distractor responses (M1: 7, M2: 2). For the remaining responses we calculated the 25^th^ percentile SRT per animal and classified all faster responses as ‘fast SRTs’. The values of these fast SRTs for target and salient distractor choices were compared with Wilcoxon rank sum tests. We also used the full range of SRTs to calculate a proportion of salient distractor choices (*p*_*SD*_ *= N*_*SD*_*/N*_*ALL*_) within a 20 ms sliding window moving through the range of SRTs with 10 ms increments.

## Supplemental figures

**Supplemental Figure 1.**
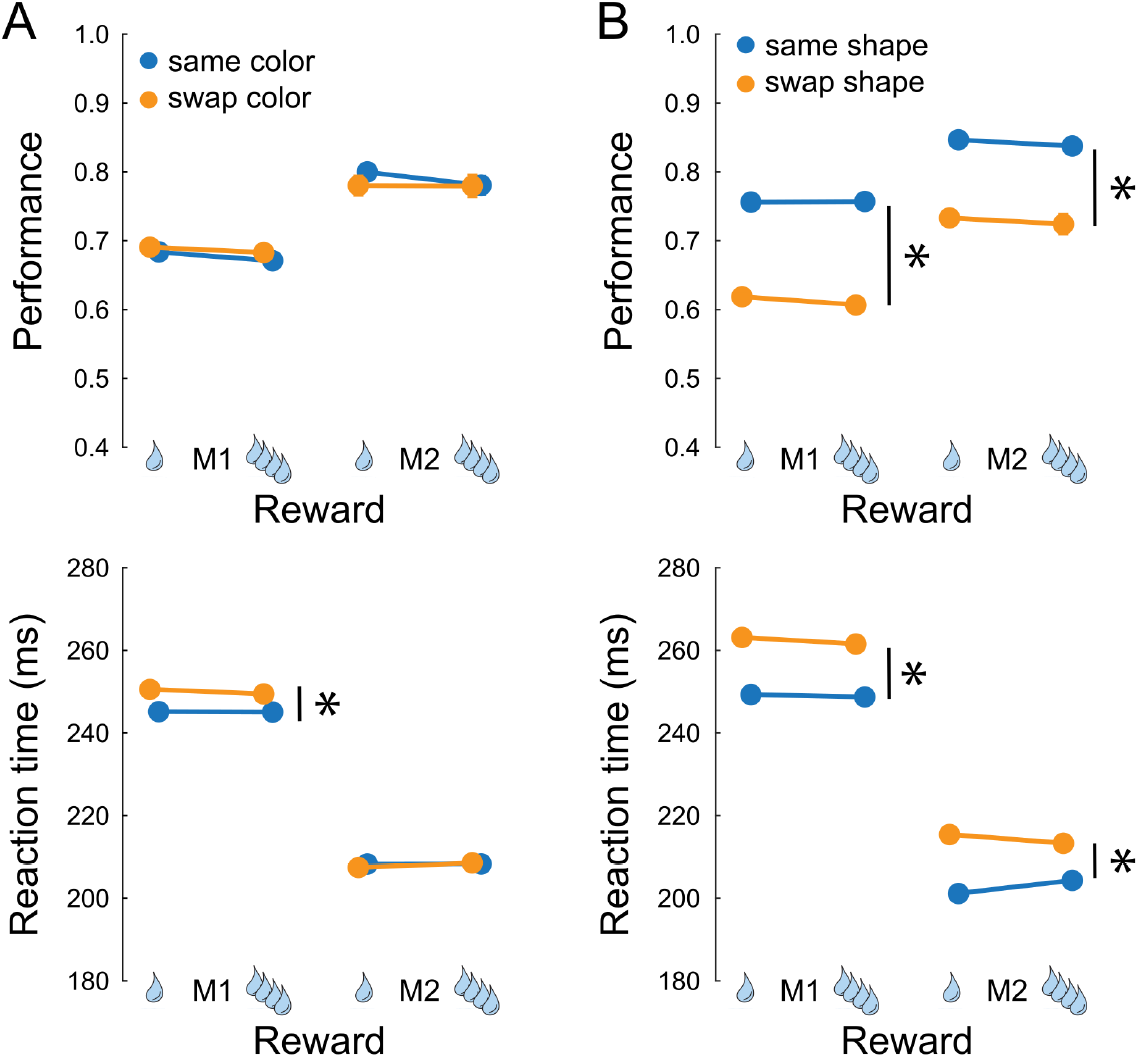
Influence of previous trial on behavioral performance. **A)** The effects of preceding reward quantity and the swapping of target and salient distractor colors (top panel) and reaction time (bottom panel) for both monkeys. Yellow lines indicate trials in which the target and distractor colors swapped relative to the previous trial; blue lines are trials in which the colors of the target and salient distractor stayed the same. Reward quantity is indicated on the horizontal axis (large rewards were four times larger than small rewards). **B)** The effects of preceding reward quantity and shape swaps on accuracy (top panel) and reaction time (bottom panel) for both monkeys. Yellow lines indicate trials in which the target and distractor shapes swapped relative to the previous trial; blue lines are trials in which the shape assignment stayed the same. Error bars (often smaller than the data points) indicate S.E.M., asterisks denote p < 0.001 for main effects as indicated by two-way ANOVAs (no interaction effects were significant at p < 0.05).

**Supplemental Figure 2.**
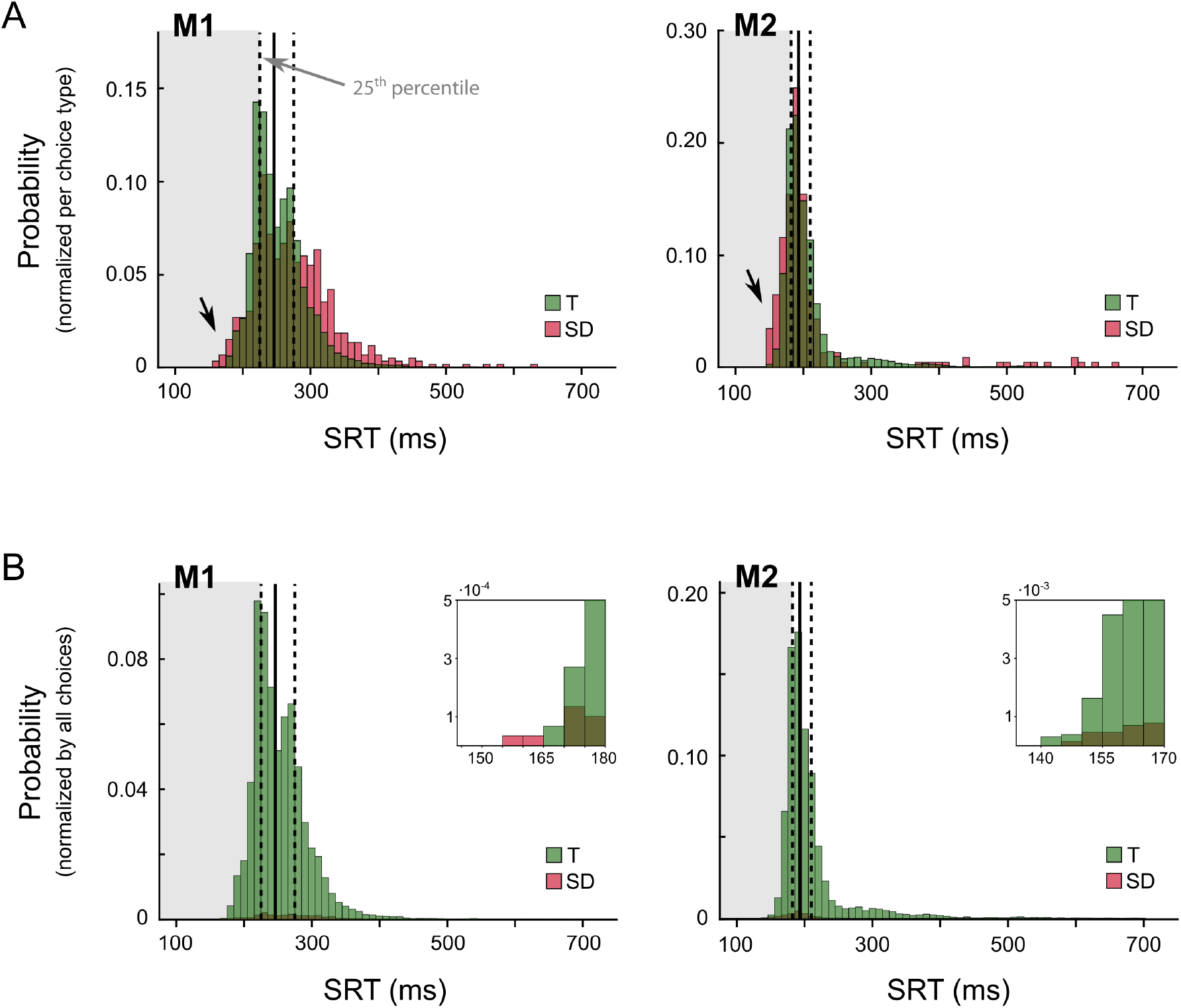
Saccadic reaction times and choices. Distributions of saccade reaction times (SRTs) for target (T, green) and salient distractor choices (SD, red) of the two monkeys (M1, M2). Solid and dashed vertical lines represent the median, 25^th^ and 75^th^ percentiles of the RT distribution. Histograms in **A)** show the distributions normalized per choice type (T or SD) as in Figure 2A, while histograms in **B)** show the distributions normalized to the total number of responses (T, SD, and ND combined). Insets in panel **B)** zoom in on the fast tail of the distributions.

**Supplemental Figure 3.**
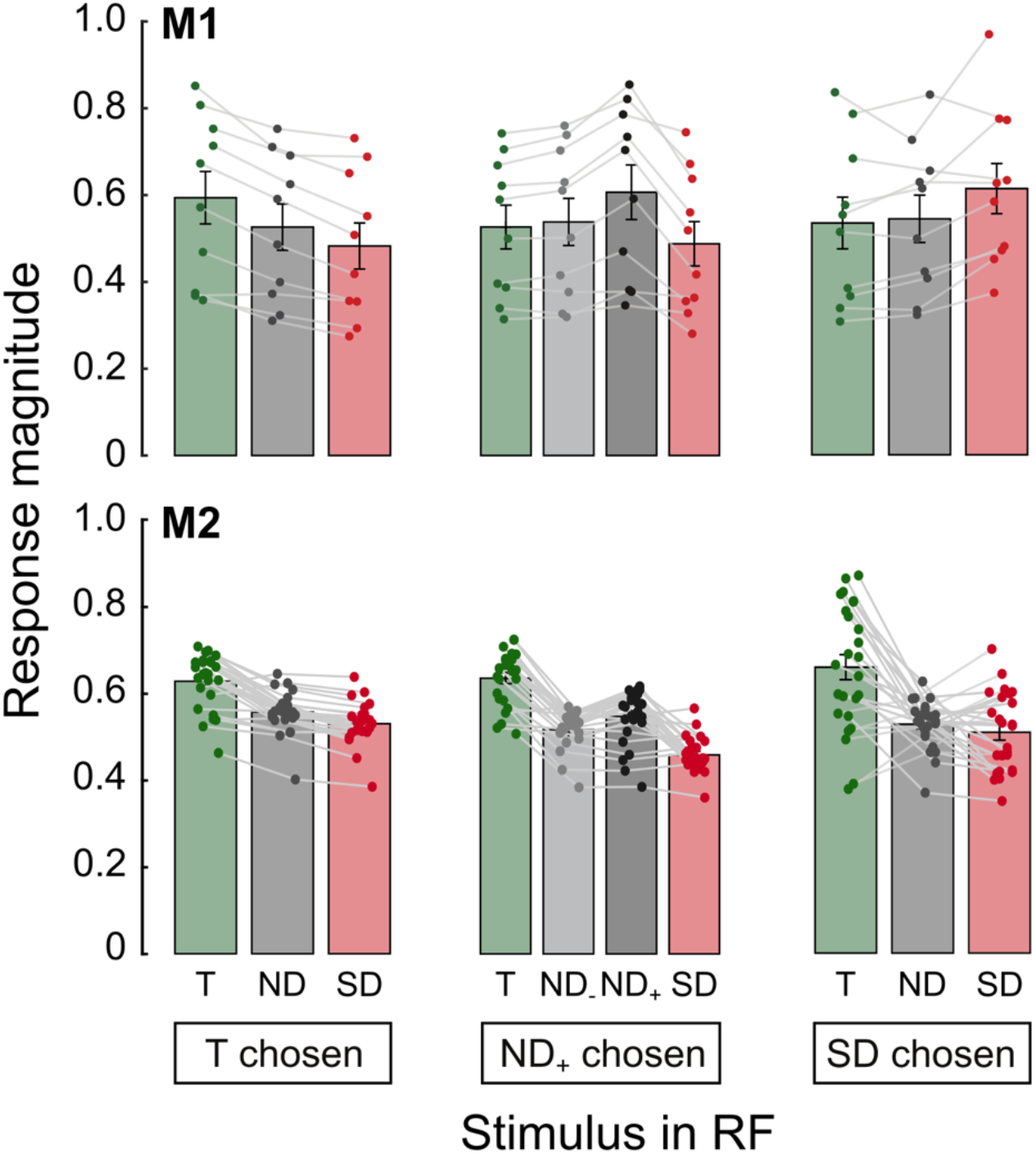
V4 activity and its dependence on choice. Average activity of V4 neurons at individual recording sites, 150-200 ms after stimulus onset. Left, correct responses to the target in M1 (top row) and M2 (bottom). Middle, erroneous responses in which the animals selected one of the non-salient distractors (ND_+_). ND_-_, response elicited by non-salient distractors that were not chosen. Right, erroneous responses in which the salient distractor was chosen. Bar colors represent the stimulus in the RF (green: target, light gray: non-salient distractor, red: salient distractor). Dark grey bars show the response elicited by the chosen non-salient distractors (ND_+_). Light grey bars show the response elicited by ND_-_. The data of individual recording sites are shown as colored data points, connected by gray lines. Error bars represent S.E.M. across recording sites.

**Supplemental Figure 4.**
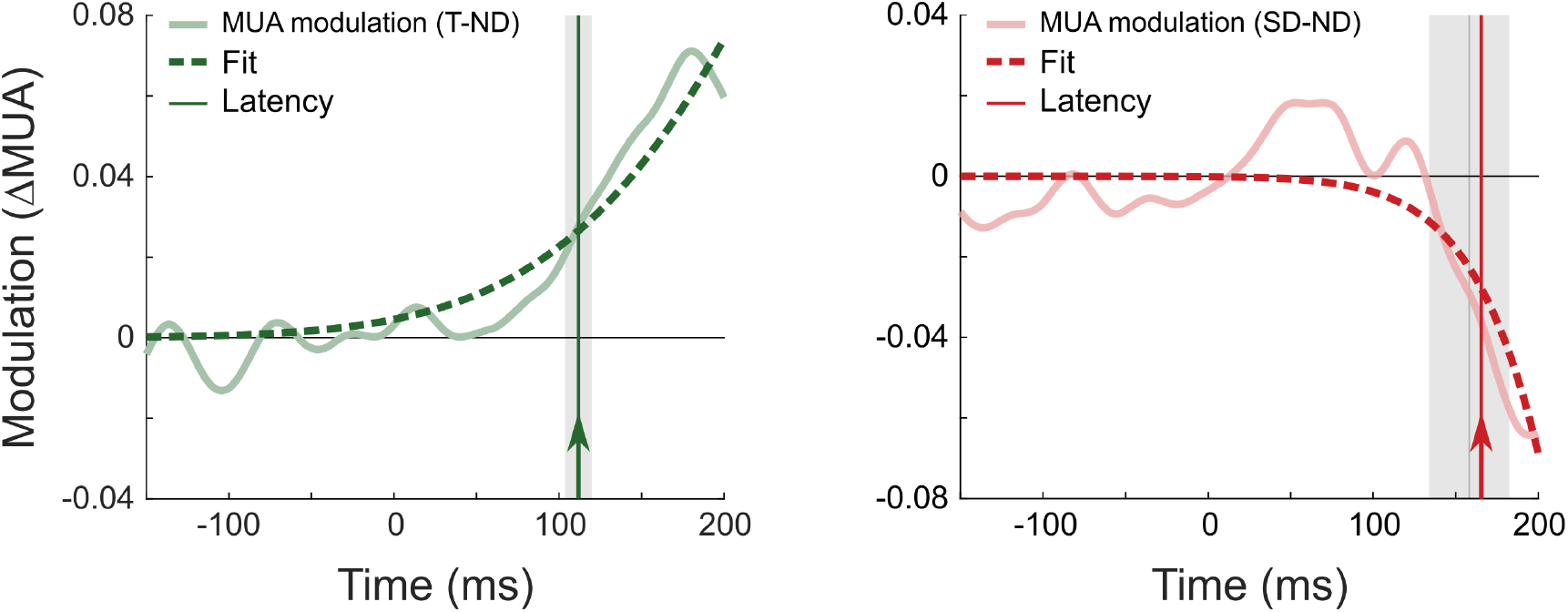
Latency analysis of target and salient distractor MUA modulation. Cumulative Gaussian functions were fit to the difference between either the average V4 activity elicited by the target and the non-salient distractor (i.e. target modulation, left panel) or the non-salient and salient distractors (i.e., salient distractor modulation, right panel). The latency (vertical line and arrow) is estimated as the time point at which the fit (dashed line) reaches 33% of its maximum. The grey area and grey vertical line indicate the mean latency and standard deviation of a bootstrap analysis (100 samples with replacement).

**Supplemental Figure 5.**
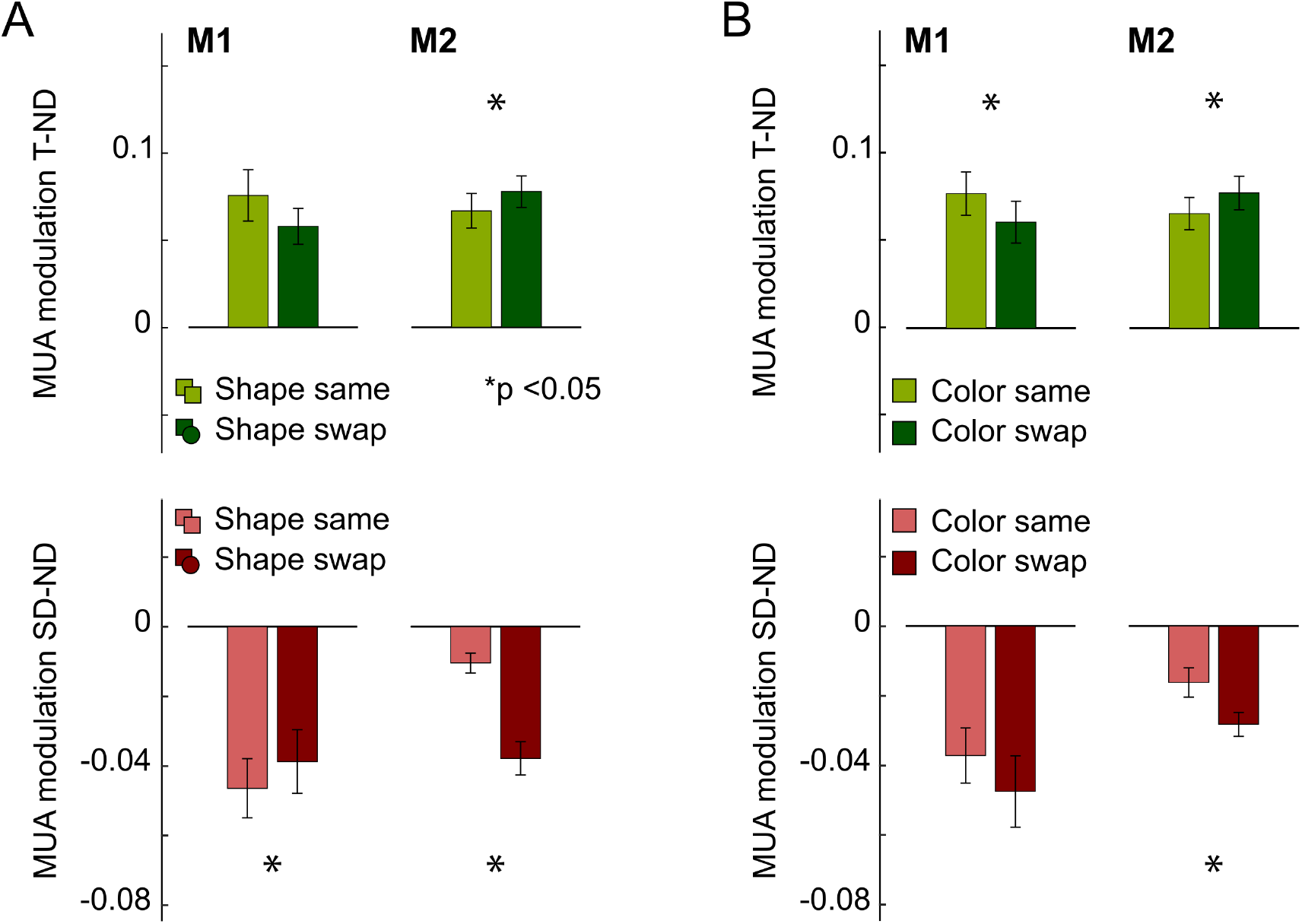
Influence of the previous trial on V4 activity. **A)** The effect of shape assignment changes on consecutive trials on the modulation of V4 activity by the target (T-ND; top row) and salient distractor (SD-ND; bottom row) in the 150-200 ms time window after stimulus onset. Bars are the mean across recording sites and error bars denote S.E.M. Light and dark colors represent trials in which the shape assignment stayed the same or changed, respectively. **B)** The effect of target and salient distractor color swapping on consecutive trials on the modulation of V4 activity by the target (T-ND; top row) and salient distractor (SD-ND; bottom row) in the 150-200 ms time window after stimulus onset. Light and dark colors represent trials in which the color assignment stayed the same or were swapped, respectively. Asterisks, significant differences (paired t-test, p < 0.05).

